# Yes Associated Transcriptional Regulator 1 (YAP1) and WW Domain Containing Transcription Regulator (WWTR1) are required for murine pregnancy initiation

**DOI:** 10.1101/2024.05.09.592984

**Authors:** Genna E. Moldovan, Noura Massri, Erin Vegter, Ivonne L Pauneto-Delgado, Gregory W. Burns, Niraj Joshi, Bin Gu, Ripla Arora, Asgerally T. Fazleabas

## Abstract

Endometrial stromal cell decidualization is required for pregnancy success. Although this process is integral to fertility, many of the intricate molecular mechanisms contributing to decidualization remain undefined. One pathway that has been implicated in endometrial stromal cell decidualization in humans *in vitro* is the Hippo signaling pathway. Two previously conducted studies showed that the effectors of the Hippo signaling pathway, YAP1 and WWTR1, were required for decidualization of primary stromal cells in culture. To investigate the *in vivo* role of YAP1 and WWTR1 in decidualization and pregnancy initiation, we generated a *Progesterone Cre* mediated partial double knockout (pdKO) of *Yap1* and *Wwtr1*. Female pdKOs exhibited subfertility, a compromised decidualization response, partial interruption in embryo transport, blunted endometrial receptivity, delayed implantation and subsequent embryonic development, and a unique transcriptional profile. Bulk mRNA sequencing revealed aberrant maternal remodeling evidenced by significant alterations in extracellular matrix proteins at 7.5 days post-coitus in pdKO dams and enrichment for terms associated with fertility-compromising diseases like pre-eclampsia and endometriosis. Our results indicate a required role for YAP1 and WWTR1 for successful mammalian uterine function and pregnancy success.

## Introduction

Successful reproduction is an impeccably complex process. The coordination of intricate molecular signals with large-scale physiological changes imparts many opportunities for error. An unfortunate statistic is that 20% of couples attempting to reproduce are infertile from either male or female contributions (NICHD). Within mammals, the fertilization of ovulated oocyte(s) is just the beginning, assuming that ovulation is not compromised. Following fertilization and early embryonic development, an embryo implants into a receptive maternal endometrium that is significantly remodeled throughout pregnancy to support gestation(1). In humans and mice, successful reproduction requires the coordinated efforts of many molecular and cellular signals and the appropriate interplay between systems, including the neuroendocrine, immune, vascular, and female reproductive tract. Humans undergo spontaneous endometrial stromal cell decidualization, a terminal differentiation of the underlying stroma within the endometrium, which is induced by increasing levels of progesterone during the secretory phase of the menstrual cycle(1). This process is critical to control trophoblast invasion and allow appropriate embryo invasion and occurs spontaneously each cycle in the absence of a conceptus(2, 3). In mice, decidualization only occurs when an embryo physically attaches to the maternal endometrial epithelium that communicates with the stroma and initiates decidualization(4). In humans, decidualization prepares the maternal endometrium for implantation while in mice implantation initiates decidualization. In both mice and humans, decidualization controls the level of trophoblast invasion and is integral to pregnancy establishment. Therefore, understanding the molecular mechanisms that underlie these complex coordinated events is essential to identifying mechanisms that may go awry and contribute to infertility.

Many molecular pathways have been implicated in decidualization and implantation, including WNT, NOTCH, and HIPPO signaling(5). Of particular interest is the mechanosensing HIPPO pathway. This pathway was first identified in *Drosophila* and so named due to mutations in this pathway leading to increased body size(6). The Hippo signaling pathway in mammals controls organ size, growth, and proliferation and does so by sensing and responding to changes in the extracellular environment, like the presence and tension of nearby cells, as well as growth factor availability(7). The Hippo pathway is a kinase cascade whereby external signals like those mentioned above induce the phosphorylation of MST1/2 through a variety of upstream signals. MST1/2 phosphorylates LATS1/2, which then phosphorylates YAP1 and WWTR1, two transcriptional cofactors. YAP1 and WWTR1 are either bound or degraded in the cytoplasm and therefore inactive when the HIPPO signaling pathway is active. Conversely, when the HIPPO signaling pathway is deactivated, YAP1 and WWTR1 translocate to the nucleus and bind their canonical partner transcription factors, the TEADs, where they regulate directly gene transcription of extracellular and cytoplasmic matrix components, cell cycle genes, and indirectly by acting as distal enhancer recruiters(7). This pathway is well conserved among mammals and has been implicated to play a role in female reproductive function.

In the murine uterus, YAP1 exhibits dynamic expression with the highest levels of mRNA and protein expression at estrus, an estrogen-dominated stage(8). In addition, YAP1 is significantly increased and phosphorylated with 17β-estradiol treatment of ovariectomized mice, suggesting YAP1’s importance in specific phases of the estrus cycle(8). Beyond the estrus cycle, YAP1 is expressed throughout the murine endometrium in early pregnancy prior to embryo implantation at embryonic day 0.5 and beyond(9). Post-implantation *Yap1* and its targets, *Ctgf* and *Ankrd1*, are significantly increased. A similar response is observed in oil-induced decidualization, suggesting a potential role for YAP1 in maternal preparation of pregnancy(9). In addition, conditional deletion of *Yap1* and its homolog, *Wwtr1* (formerly known as *Taz*), under *Anti-Mullerian hormone receptor type 2* driven *Cre* expression, results in degradation of the oviductal isthmus myosalpinx complicating embryo transport, suggesting an essential role for YAP1 and WWTR1 in structural integrity(10). Beyond what is known in the mouse, YAP1 and WWTR1 have been independently identified as critical for endometrial stromal cell decidualization *in vitro*. YAP1 expression increases in the first 48 hours of *in vitro* endometrial stromal cell decidualization, and knockdown by shYAP prior to induction of *in vitro* decidualization results in a compromised decidualization response(11). In addition, WWTR1 increases during in vitro decidualization at day 6, and the knockdown of WWTR1 also compromises the expression of decidualization markers IGFBP1 and dPRL (unpublished)(12). These studies highlight the roles of YAP1 and WWTR1 in *in vitro* decidualization, but this potential has yet to be explored *in vivo*. In this study, we generated conditional knockout of YAP1 and WWTR1 under the *Progesterone receptor Cre* to explore the potential roles and regulation of these Hippo homologs in pregnancy.

## Results

### Generation of Yap1 and Wwtr1 partial double knockout

*Progesterone (Pgr) Cre* expressing male mice were crossed to double floxed females (*Yap^f/f^ Wwtr1^f/f^*) to generate heterozygous founders. Offspring were then backcrossed to generate *Pgr^Cre/+^ Yap^f/f^ Wwtr1^f/f^* females for fertility investigation. The resulting mice were genotyped as *Pgr^Cre/+^ Yap^f/+^ Wwtr1^f/f^* (pdKO) or *Pgr^+/+^ Yap^f/f^ Wwtr1^f/f^* but never as *Pgr^Cre/+^ Yap^f/f^ Wwtr1^f/f^*. Investigation into the Mouse Genome Informatics database revealed that *Progesterone* and *Yap1* are approximately 900kB or 0cM apart, and therefore, it was impossible to generate a full double knockout model utilizing the *Pgr Cre*. The investigation continued into the role of *Yap* and *Wwtr1* in the murine reproductive tract utilizing the partial double knockout model.

### Progesterone Cre conditional deletion of Yap1 and Wwtr1 exon 3 does not affect reproductive tract structure

Uteri and ovaries were assessed for Cre-mediated recombination of the *Yap1* and *Wwtr1* alleles (Figure 1). At 3.5dpc, neither *Yap1* nor *Wwtr1* were differentially expressed in uteri utilizing primers that amplify the floxed exon 3, but the target gene *Ccn2* was significantly decreased (unpaired t-test *p*=0.032, Figure 1A-D). However, at 5.5dpc, *Wwtr1* was significantly decreased in pdKO uteri (unpaired t-test *p*=0.017, Figure 1B). Immunohistochemical analysis indicated epithelium and stromal compartment-specific expression of YAP1 and WWTR1 in uteri at 3.5dpc, however expression levels were not significantly different between controls and pdKOs (Figure 1E-F). Evidence of *Yap1* and *Wwtr1* and target gene depletion was not evident in whole ovaries (Supplementary Figure 1A). In addition, both YAP1 and WWTR1 protein expression was not significantly different in granulosa cells of 3.5dpc control and pdKO ovaries (Supplementary Figure 1B). In addition to these molecular analyses, no structural differences were observed in the ovaries, oviducts, and uteri of virgin females at proestrus or 3.5dpc (Supplementary Figure 2A-B).

**Figure 1.**
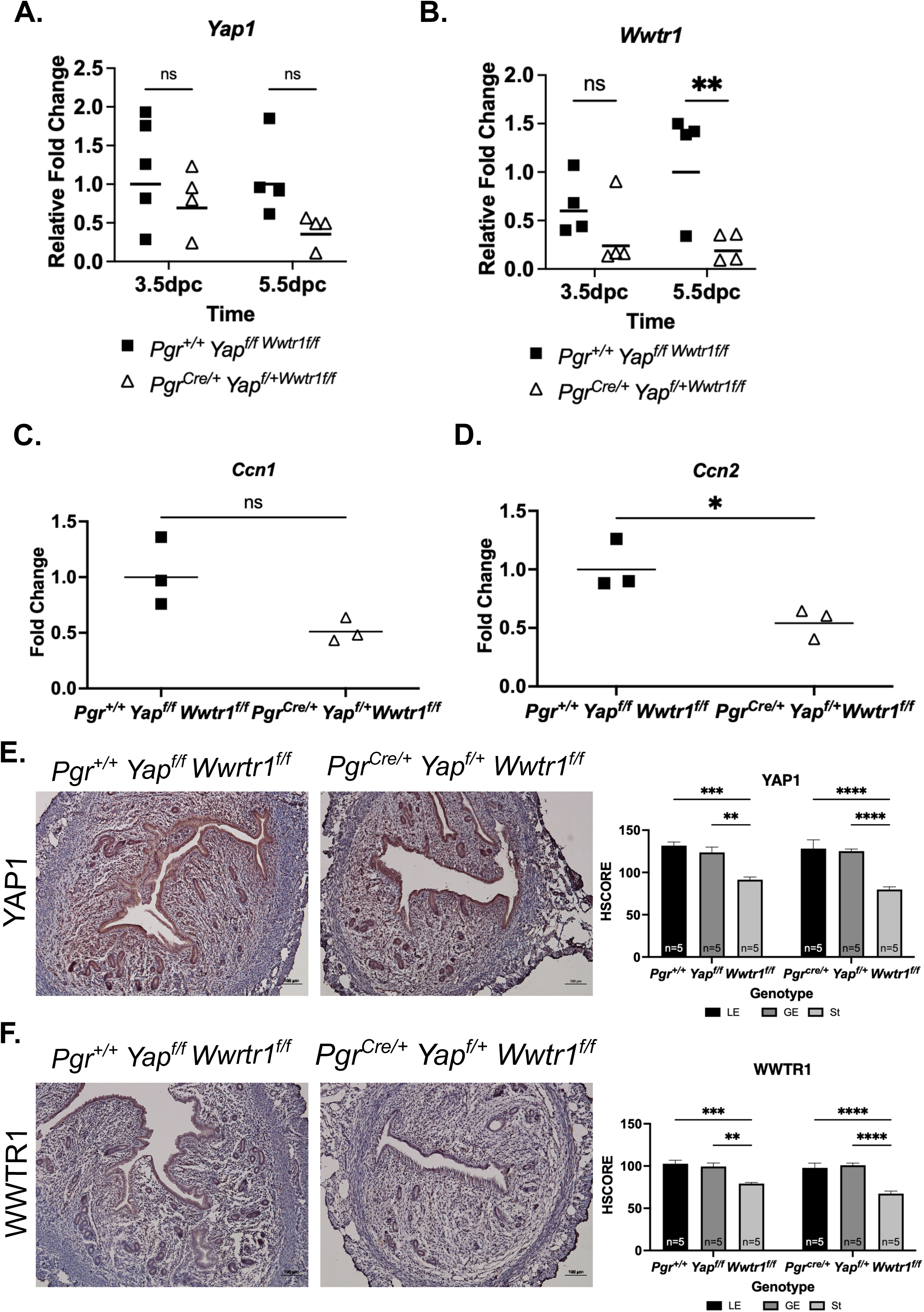
YAP1 and WWTR1 expression at 3.5 and 5.5dpc. **A.** *Yap1* mRNA expression is not significantly different in pdKOs compared to controls in early pregnancy. **B.** *Wwtr1* mRNA expression is significantly decreased in pdKOs at 5.5dpc but not at 3.5dpc compared to controls. **C.** YAP/WWTR1 target gene *Ccn1* is not differentially expressed at 3.5dpc in pdKOs. **D.** *Ccn2*, a YAP/WWTR1 target gene, is significantly decreased at 3.5dpc in pdKOs. **E.** YAP1 protein expression at 3.5dpc and semi-quantitative HSCORE. **F.** WWTR1 protein expression at 3.5dpc and semi-quantitative HSCORE.

### Partial ablation of Yap and Wwtr1 induces subfertility

*Pgr^Cre/+^ Yap^fl/+^ Wwtr1^fl/fl^* partial double knockouts (pdKO) produced significantly fewer total pups throughout a 6-month breeding trial (n=4, Mixed-effects analysis factor genotype, *p*=0.006, Figure 2A). In addition, pdKO females had smaller litter sizes than controls (unpaired t-test *p*=0.0007, Figure 2B). The pdKO mice bore approximately 50% fewer pups and litters than the floxed controls throughout the breeding trial (Supplementary Table 3). In addition, the pdKO mice displayed a reduced capacity to produce pups as time progressed (Mantel-Cox test *p=*0.007, Figure 2C). One individual only produced two live-born litters when placed with multiple fertile males (Figure 2D). In addition to the subfertility shown throughout the breeding trial, pdKO females in an expanded cohort exhibited significantly reduced first and second litter sizes (Mixed effects analysis factor genotype, *p*=0.0026, Supplementary Figure 3A). This was also associated with increased inter-litter timing (Mixed effects analysis factor genotype *p*<0.0001, factor litter number *p*=0.0032, Supplementary Figure 3B). Despite increased inter-litter timing, pdKO body weight measured over an 8-week period matched that of the wild types (Supplementary Figure 3C). At the termination of the 6-month breeding trial, n=3 mice per genotype were collected for analyses. These mice did not display any significant differences in body, uterine, or ovarian weight at the time of collection (not shown), and no alterations in reproductive tract structure were observed grossly nor with histological analyses (Supplementary Figure 2C).

**Figure 2.**
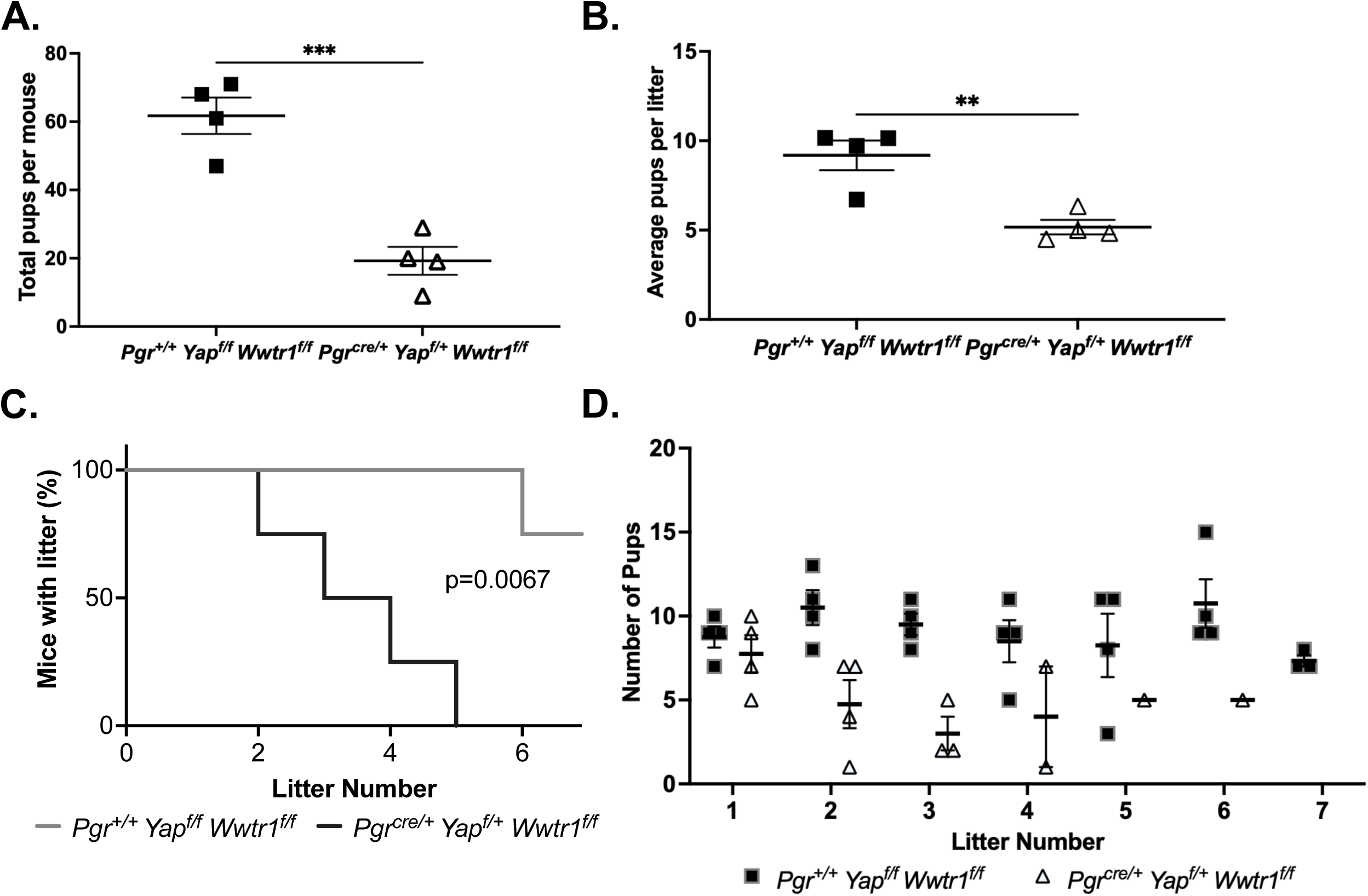
pdKOs are subfertile with smaller litter sizes. **A.** pdKO females had fewer total pups across a 6-month breeding trial. **B.** pdKO females had smaller litter sizes compared to controls across a 6-month breeding trial. **C.** pdKO females exhibited a reduced capacity to produce pups over time. **D.** pdKO females began to lose the ability to bear live born litters after the second litter.

### Yap1-Wwtr1 partial double knockout females exhibit decreased decidualization response

Artificial decidualization was maintained for 5 days, and then uteri were collected for analysis. Representative images show unstimulated (left) and stimulated (right) horns for floxed controls and pdKOs (Figure 3A). The uterine wet weight ratio of stimulated to unstimulated horn was not different in the pdKO group compared to controls (Figure 3B). However, mRNA expression of decidualization response genes, *Wnt4* and *Bmp2*, indicated a blunted decidualization response in the stimulated uterine horns of pdKOs compared to controls (Two-way ANOVA *p*=0.04 *Wnt4*, and *p*=0.006 *Bmp2*, Figure 3C).

**Figure 3.**
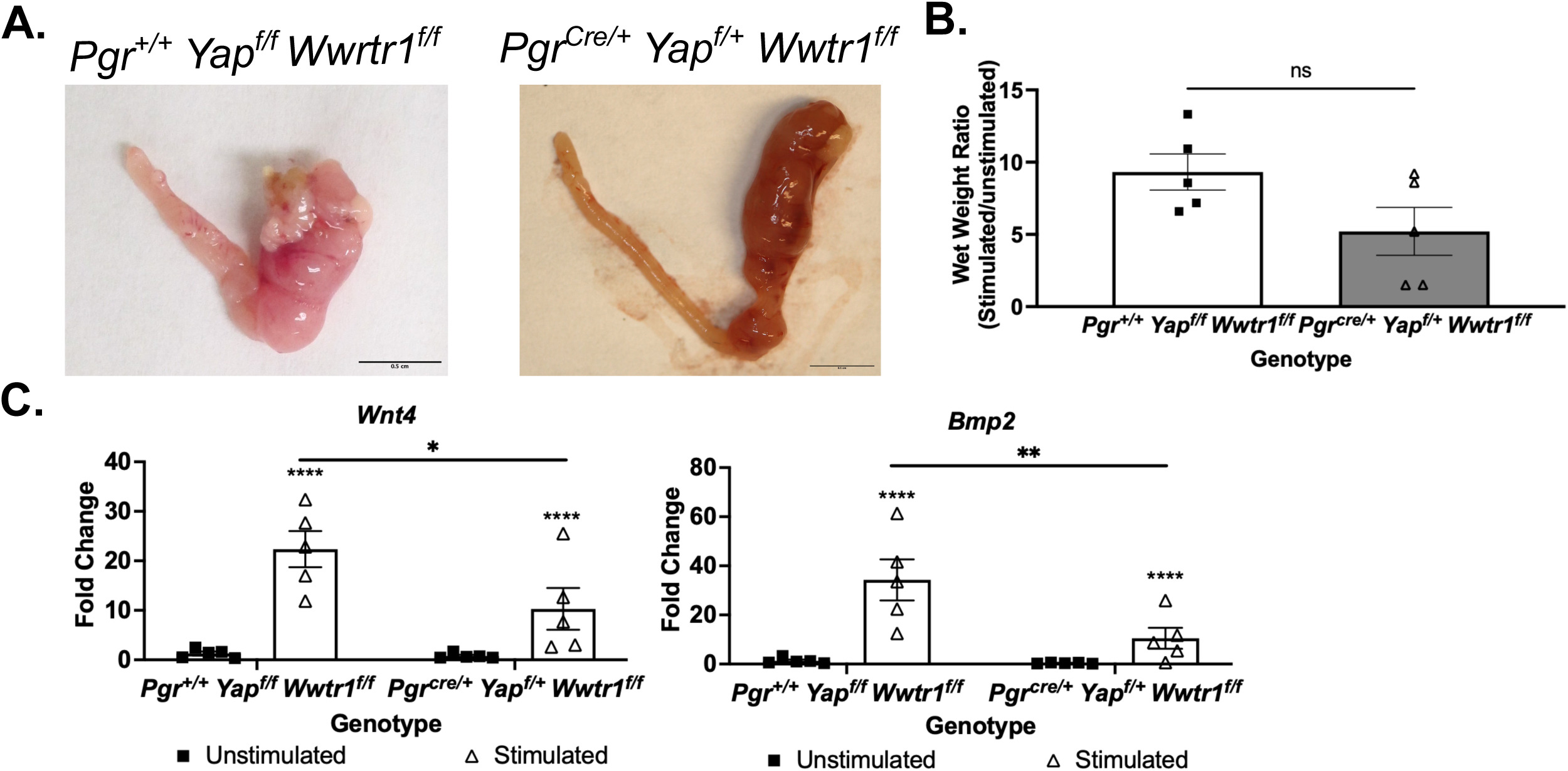
Loss of *Yap1* and *Wwtr1* compromises the decidualization response. **A.** Representative micrographs of unstimulated (left) and stimulated (right) uterine horns 5 days after artificial decidualization induction. **B.** Uterine wet weight ratio of stimulated over unstimulated horn after 5 days of artificial decidualization. **C.** mRNA fold change of decidualization markers *Wnt4* and *Bmp2* 5 days after artificial decidualization induction.

### Loss of Yap1 and Wwtr1 leads to incomplete pregnancy failure

Pregnancy was assessed in control and pdKO mice at various time points including implantation (4.5dpc, n=6-8), post-implantation (5.5dpc, n=6-10), early pregnancy (7.5dpc, n=5-7), placentation initiation (9.5dpc, n=5-7), and post-placentation (12.5dpc, n=8-10) (Figure 4A-B). The gross morphology of the uteri was normal at all time points except 12.5dpc when pdKO mice exhibited many resorption sites (Figure 4A). Quantification of normal implantation sites, when assessed by blue dye reaction, revealed variance in phenotype. At 4.5dpc, the time of implantation, pdKO females were split with 50% of mice with positive implantation sites with blue dye injection and 50% of mice without positive implantation sites (Figure 4B). However, by 5.5dpc, most pdKO females had positive implantation sites with numbers comparable to controls (Figure 4B). At 7.5dpc and 9.5dpc, the number of mice with positive implantation sites decreased to ∼40%, with ∼60% not having positive implantation sites at 9.5dpc (Figure 4B). At 12.5dpc, embryonic loss was evident with visually identifiable resorption sites (Figure 4B). In addition, across all time points measured, pdKO females exhibited a higher likelihood of being not pregnant compared to controls (33% of individuals, Fisher’s exact test *p*<0.0001, Figure 4C).

**Figure 4.**
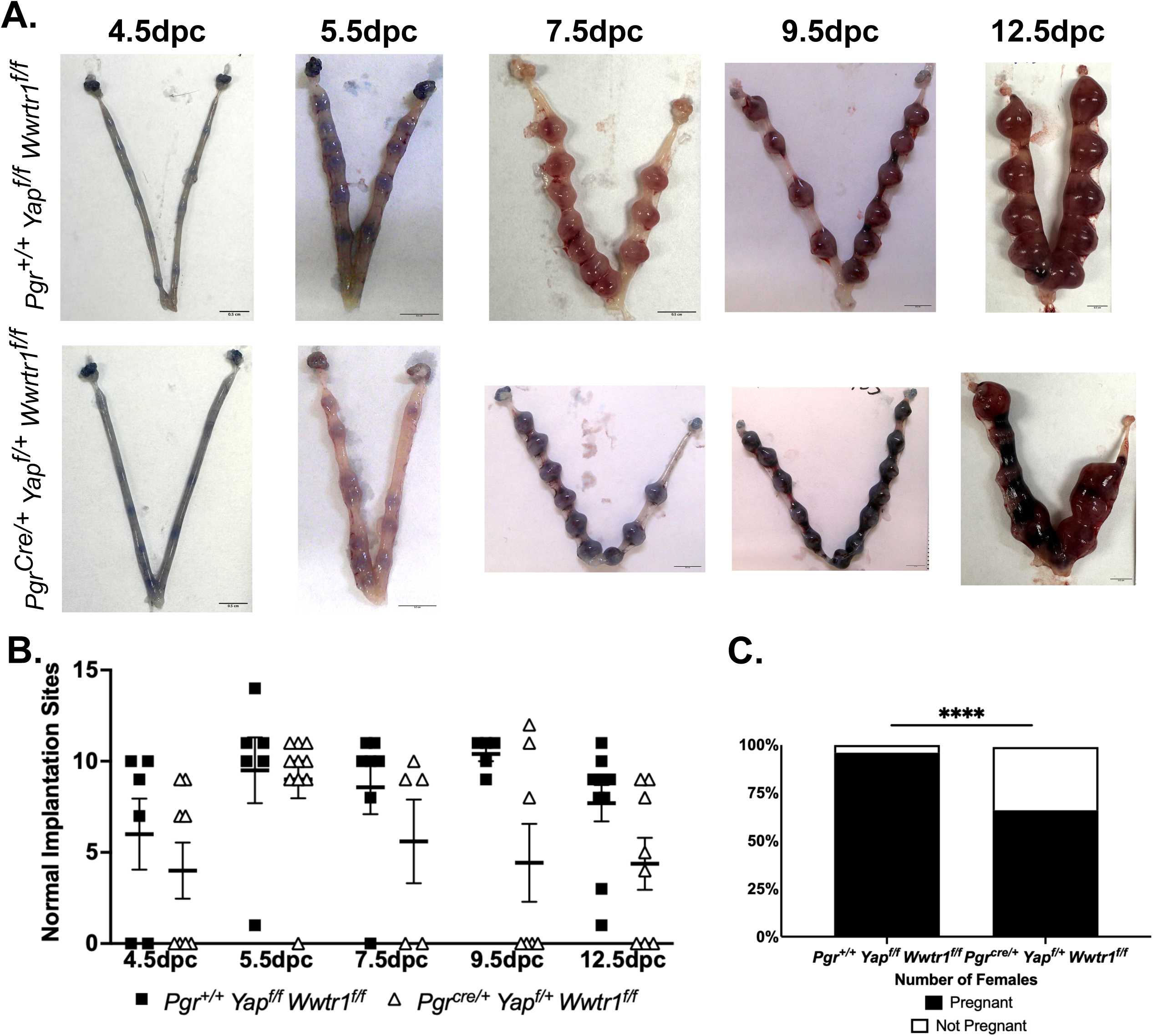
pdKOs exhibit fetal loss at 12.5dpc. **A.** Representative micrographs of control (top panel) and pdKO (bottom panel) uteri at various stages of pregnancy. Scale bars = 0.5cm. **B.** Quantification of normal implantation sites across pregnancy in both control and pdKO mice (Mixed-effects model comparison genotype *p*=0.0154). **C.** pdKO females exhibit a higher rate of nonpregnancy compared to controls across all timepoints collected (Fisher’s Exact Test, *p*=0.0058).

### pdKOs display modest interruptions in embryo transport

Determination of the cause of pregnancy failure prompted the investigation into ovulation and fertilization rates and embryo transport. Post-fertilization at 1.5dpc, when all ovulated oocytes should be in the oviducts, we noted comparable numbers of unfertilized oocytes (UFO) and 2-cell embryos in pdKOs and controls (Figure 5A). However, at 3.5dpc, the time of embryo entry into the uterus, we noted the emergence of two mutually exclusive groups: pdKO females with embryos found only within the oviducts and pdKO females with embryos found only within the uterus (Figure 5B-C). The total number of visual corpus lutea did not differ between pdKOs and controls, nor did the total number of embryos flushed at 3.5dpc (Figure 5D). In addition, visualization of oviducts indicated appropriate folding patterns (Supplementary Figure 4A)(13). Individual uteri were imaged utilizing tissue clearing to allow visualization of embryos using the whole-mount immunofluorescent protocol. Embryos found within pdKO uteri at 3.5dpc were 50% of the quantity found within control uteri (Supplementary Figure 4B and 4C).

**Figure 5.**
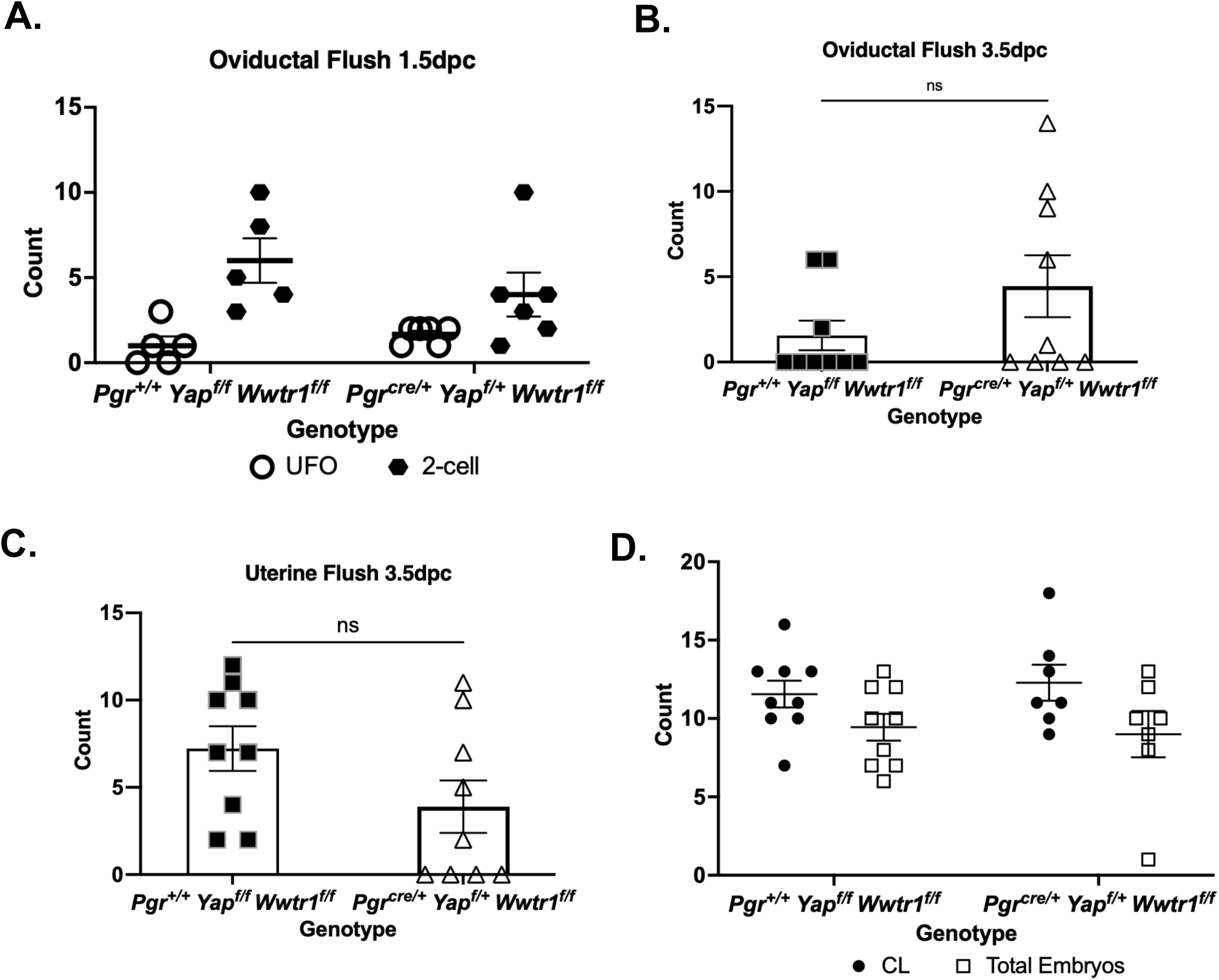
pdKOs females have delayed uterine embryo entry. **A.** Quantity of unfertilized oocytes (UFO) and 2-cell embryos flushed from oviducts at 1.5dpc from floxed controls and pdKOs. **B.** Total embryos flushed from oviducts of floxed controls and pdKOs at 3.5dpc. **C.** Total embryos flushed from uteri of floxed controls and pdKOs at 3.5dpc. **D.** Total corpus lutea (CL) and embryos flushed at 3.5dpc.

### pdKOs exhibit altered maternal endometrial receptivity

To disseminate the cause of delayed embryo entry into the maternal endometrium combined with partially delayed implantation and to rationalize delayed embryonic development, we investigated endometrial receptivity at 3.5dpc (Figure 6). The progesterone receptor expression by both mRNA and protein was not altered in pdKO uteri at 3.5dpc (Figure 6A and B). However, we noted significantly decreased target gene expression, *Ihh,* and *Nr2f2*, in 3.5dpc uteri (unpaired t-test, *p*<0.01, Figure 6A). In addition, luminal epithelial expression of proliferation marker Ki67 was maintained in pdKO uteri (Figure 6C). Interestingly, estrogen receptor signaling decreased in pdKOs compared to controls at 3.5dpc (Unpaired t-test, *p*<0.03, Supplementary Figure 5).

**Figure 6.**
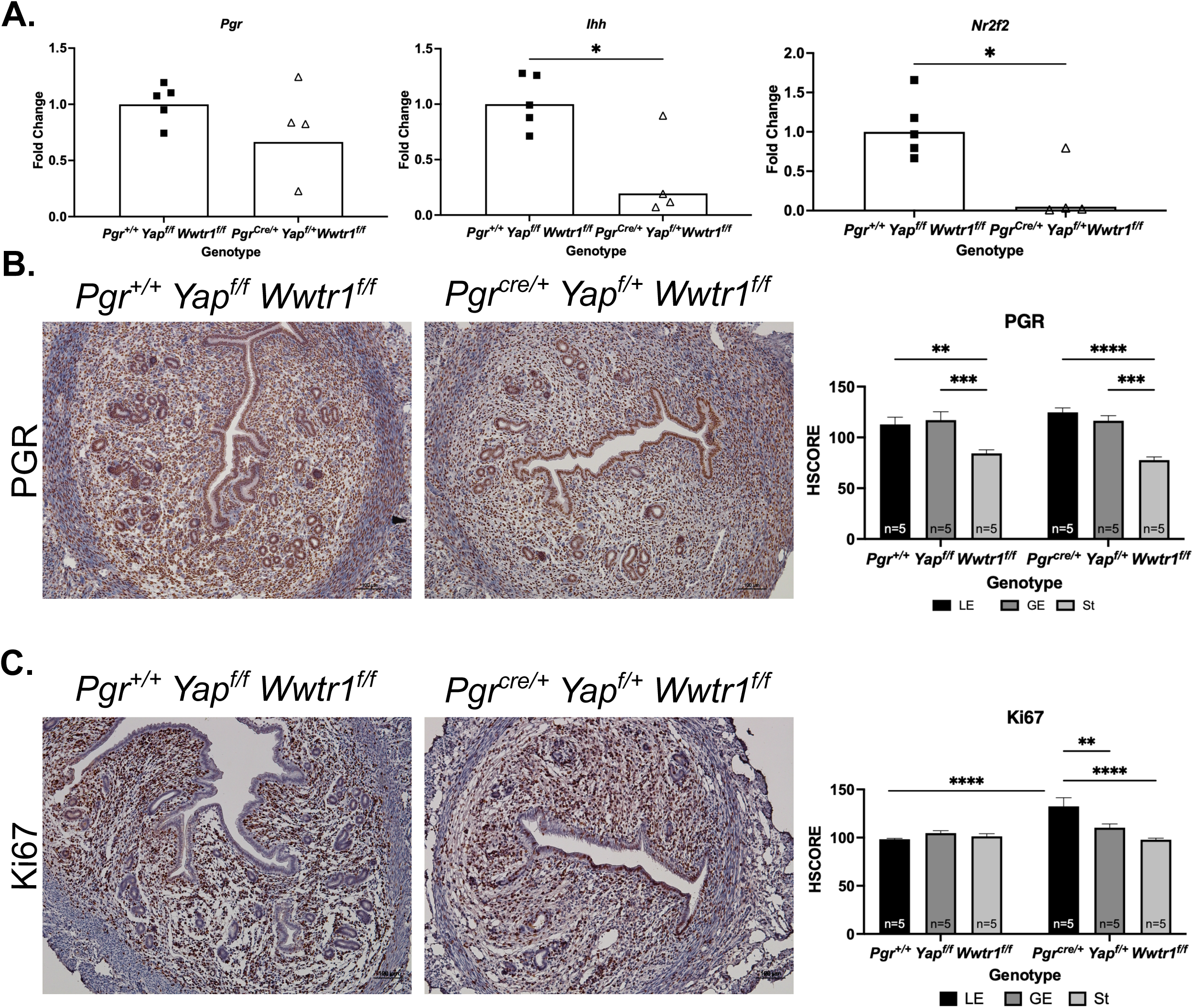
*Yap1* and *Wwtr1* are required for appropriate endometrial receptivity. **A.** mRNA relative fold change of *Pgr* and two target genes *Ihh* and *Nr2f2* at 3.5dpc in whole uteri of controls and pdKOs. **B.** Immunohistochemical staining for PGR and semi-quantitative H-CORE of DAB signal in control and pdKO uteri at 3.5dpc. **C.** Immunohistochemical staining for Ki67 and semiquantitative HSCORE of DAB signal at 3.5dpc in control and pdKO uteri.

### pdKO uteri contain embryos exhibiting delayed development

Implantation chamber morphology was notably normal when visualized with whole uterine imaging at 5.5dpc (Figure 7A). However, pdKO uteri contained embryos that morphologically appeared delayed with decreased elongation compared to controls at 5.5dpc (Figure 7B). Most embryos visualized (59%) within pdKO uteri were delayed (Figure 7C). Overall, implantation chamber length was not affected despite delayed embryos being found in most implantation sites within pdKOs visualized (Figure 7D). Delayed embryos with decreased elongated morphology had 3D surface volumes significantly lower compared to normal elongated embryos found within both control and pdKO implantation sites (unpaired t-test, *p=*0.01 compared to normal control embryos and *p*=0.001 compared to normal pdKO embryos Figure 7E). Additionally, the ratio of embryo volume to implantation chamber length of those embryos that were identified as delayed was significantly decreased (unpaired t-test, *p*<0.01, Figure 7F). This finding necessitates further investigation of the factors contributing to delayed embryo development and implantation in our mouse model.

**Figure 7.**
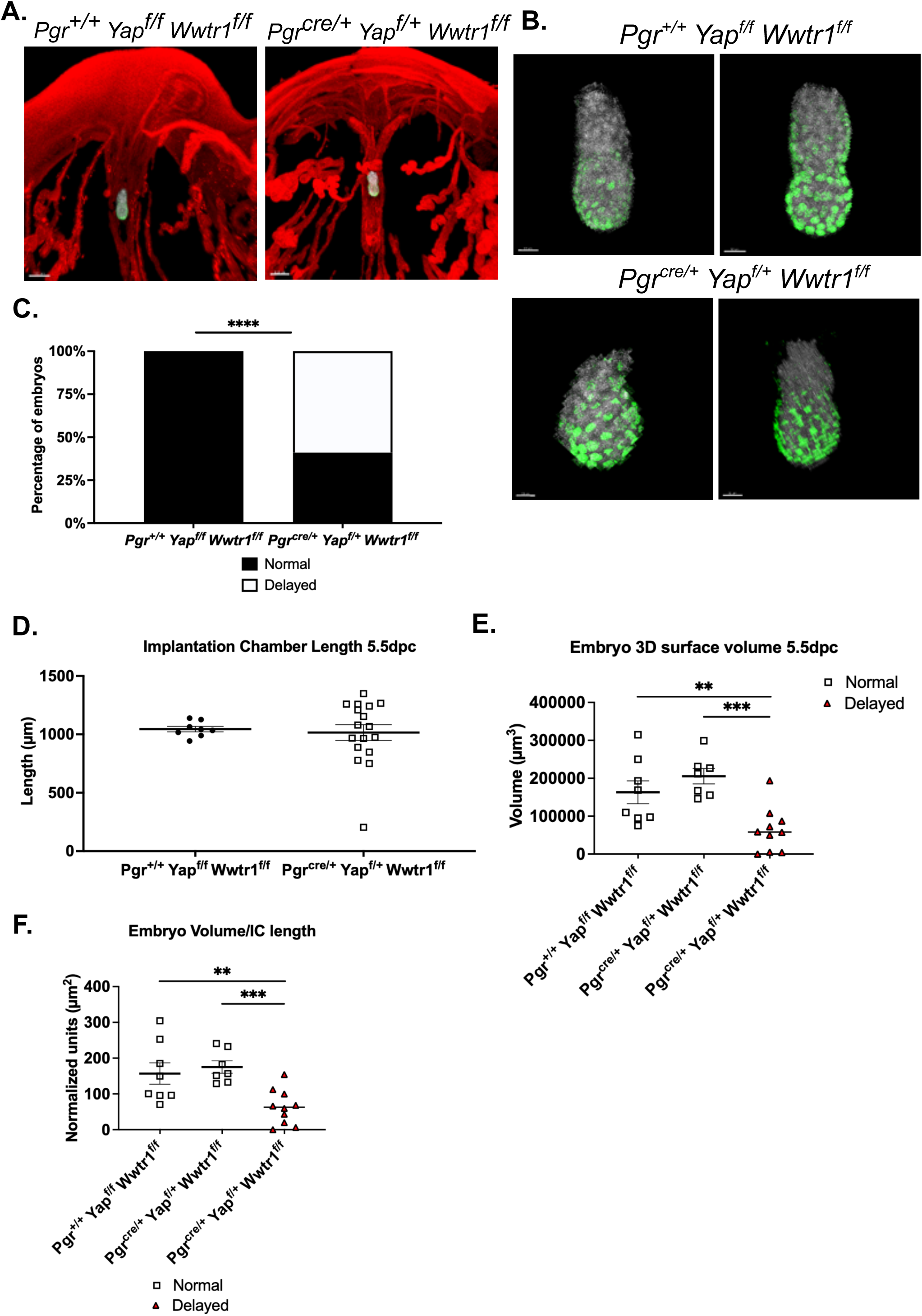
Loss of *Yap1* and *Wwtr1* in maternal endometrium induces delayed embryonic development. **A.** Representative micrographs of whole mount uterine imaging of individual implantation chambers and embryos at 5.5dpc (red=E-cadherin, white=Hoechst, Green=FOXA2). **B.** Representative embryos from controls and pdKOs at 5.5dpc (white=Hoechst, Green=FOXA2). **C.** Quantification of normal and delayed embryos imaged at 5.5dpc. **D.** Plotted implantation chamber length for controls and pdKOs at 5.5dpc. **E.** 3D embryo surface volume of normal and delayed embryos at 5.5dpc. **F.** Ratio of embryo volume to respective implantation chamber length for normal and delayed embryos at 5.5dpc.

### pdKOs have a unique transcriptional profile

Whole uterine cross-sections at 7.5dpc from pdKO (n=3) and floxed controls (n=4) were subjected to bulk mRNA sequencing to elucidate the transcriptional repercussions associated with the loss of *Yap1* and *Wwtr1* in the pregnant uterus. Samples were separated on an MDS plot based on genotype (Figure 8A). We identified 16,674 total genes after filtering and 1,785 differentially expressed genes (DEG), with 884 genes upregulated and 901 downregulated in the knockouts compared to controls (Figure 8B). Differentially expressed genes (DEG) included eight genes from the Hippo signaling pathway including upstream regulators (*Amotl1* and *Amotl2*), kinases (*Sav1* and *Lats2*), transcription factor (*Tead4*), transcriptional effectors *(Yap1* and *Wwtr1*), and target genes (*Ccn2* and *Birc5*) (Figure 8C). Decreased differentially expressed genes also included four decidualization response genes (Figure 8D). DEG contributed to Gene Ontology Cellular Component terms like extracellular matrix, collagen trimer, and collagen-containing extracellular matrix, indicating that loss of *Yap1* and *Wwtr1* affected genes involved in cellular structure (Figure 8E). In addition, upregulated genes were enriched for Rat Genome Database (RGD) disease terms like pre-eclampsia, endometriosis, liver cirrhosis, and pulmonary fibrosis, indicating that pdKO uteri were enriched for genes connected with reproductive diseases associated with pregnancy loss and fibrosis, which has classically been associated with Hippo signaling (Figure 8F).

**Figure 8.**
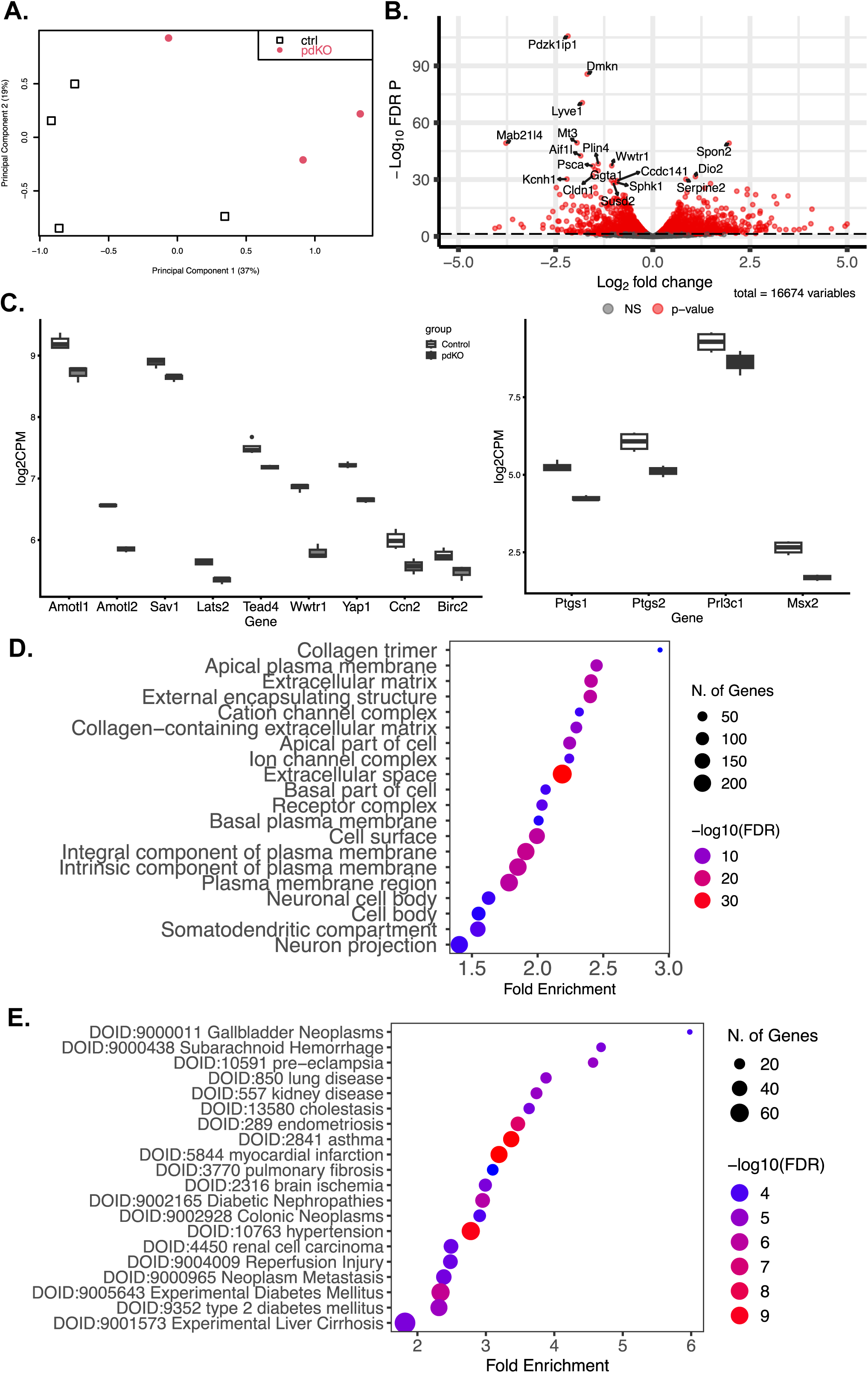
Transcriptional changes associated with *Yap1* and *Wwtr1* function during pregnancy. **A.** MDS plot of all uteri samples sequenced at 7.5dpc for controls and pdKOs. **B.** Volcano plot of all genes identified with significantly differentially expressed genes shown in red. **C.** Log2CPM values of significantly differentially expressed Hippo signaling genes and decidualization associated genes in pdKO uteri compared to controls. **D.** Normalized exon 3 (floxed or excised exon) counts for *Yap1* and *Wwtr1* in control and pdKO uteri. **E.** GO cellular component enrichment for all DEG. **F.** Enrichment of upregulated DEG for Disease RGD terms.

**Figure 9.**
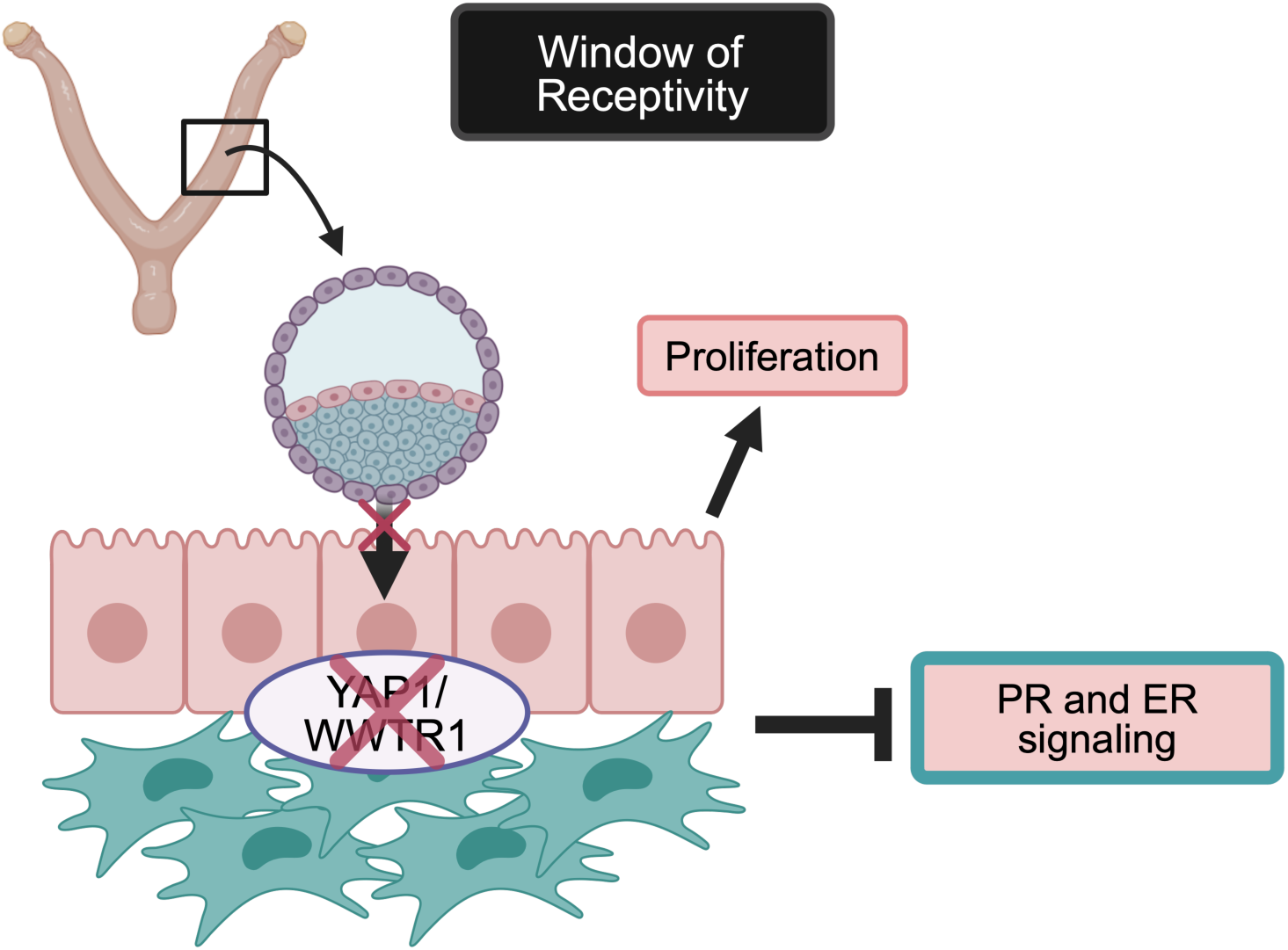
Summary Figure. YAP and WWTR1 are required for Progesterone and Estrogen receptor (PR and ER) signaling during the window of receptivity. Blastocyst attachment is delayed likely due to maintained proliferation in the luminal epithelium. In addition, loss of YAP1 and WWTR1 in both the endometrial epithelium and stroma leads to suppressed ER and PR signaling compromising pregnancy initiation.

## Discussion

The deficient uterine receptivity and dysregulation of hormonal signals within the pdKO females is interesting. However, the interplay between the Hippo signaling pathway and the hormonal response has been observed before. The functional kinases of the Hippo kinase cascade, serine/threonine kinase 3 and 4 (Stk3/4), have been shown to be differentially expressed throughout the murine estrous cycle primarily under the control of estrogen signaling(14). Indeed, YAP1 expression is also differentially regulated throughout the murine estrous cycle, with its level being highest during estrous and diestrus(8). Importantly, estrus in female mice is when ovulation occurs, and 3 days later is when the window of receptivity begins suggesting that YAP1 is present and may play a role in hormone response during this period. In addition to the cyclical presence of YAP1 as well as activation of the Hippo signaling pathway, the function of this pathway has also been associated with hormonal action. In disease states like breast cancer and endometriosis, YAP1 bound to TEAD transcription factors can induce transcription of *ESR1,* or alternatively, in endometriosis, YAP1 can regulate upstream factors that influence *PGR* expression (15, 16). These reports support an indirect method of hormonal control and interpretation of the signal, but it is possible within the canonical function of YAP1/WWTR1 as transcriptional coactivators that they could play a direct role in hormone receptor function. In one study, YAP/TEAD complexes bound to ESR1 enhancer regions and estrogen response elements (17). We propose that within our model, it is possible that the loss of *Yap1/Wwtr1* and subsequent binding to TEADs, prevents the acquisition and activation of distal estrogen and progesterone receptor response elements and enhancer recruiters. This would explain why the loss of *Yap1/Wwtr1* during the receptive window leads to a failure of PGR and ESR1-regulated genes to be activated despite appropriate expression of receptor levels during this time (Summary Figure). The mechanistic aspect of this hypothesis remains to be tested but opens a path for novel roles of YAP1/WWTR1 function within the murine uterus.

We showed that depletion of *Yap1* and *Wwtr1* led to a lack of decidualization response and maternal remodeling. YAP1 and WWTR1 are classically noted to respond to extracellular changes in stiffness transduced through the Hippo signaling pathway and decidualization and pregnancy processes, which rely on appropriate extracellular changes to alter the maternal endometrial environment(7). They also classically regulate the transcription of extracellular and structural component genes in addition to cell cycle regulatory genes(7). Unsurprisingly, the loss of Yap1/Wwtr1 led to the enrichment of DEG in GO Cellular Component terms like plasma membrane, collagen-containing matrix, and extracellular matrix (Supplementary Table 6). Within the upregulated and downregulated DEG of the bulk mRNA sequencing, specific classes of genes included extracellular matrix genes such as collagens, CCNs, laminins, and extracellular remodeling genes like matrix metalloproteases (Supplementary Tables 6 and 7). However, differential gene expression analysis of whole tissue makes it difficult to determine whether altered wound healing or fibrosis occurs in the pdKO females. Given the phenotypic data that show a progressive loss of fertility in the pdKO females, it is likely that either aberrant wound repair mechanisms or increased scarring and associated fibrosis are occurring. Both wound healing and fibrosis-associated markers were found within the DEG at 7.5dpc, including the decrease of angiogenic markers *Vegfc*, *Vegfb*, increased *Pdgfrb*, and increased expression of *Il13ra1* and *Il4ra1*. Angiogenesis is associated with wound healing but is negatively associated with fibrosis(18). These results suggest that loss of *Yap1*/*Wwtr1* leads to decreased wound healing during pregnancy through the downregulation of proangiogenic factors. However, vasculature responds to a lack of angiogenic signal by upregulating receptor expression on the vascular smooth muscle(18). In addition, IL13RA1 forms a heteromeric complex with IL4RA and activation of this receptor complex is widely considered pro-fibrotic, supporting that the pdKO uteri may be exhibiting increased fibrosis(18). Indeed, pathway enrichment for upregulated DEG in Disease RGD included disease terms for fibrotic diseases like pulmonary fibrosis and liver cirrhosis (Supplementary Table). It is likely that loss of *Yap1/Wwtr1* contributes to both a lack of wound healing and increased fibrosis within the murine uterus during pregnancy and this contributes to the repetitive pregnancy loss seen within the pdKO females.

The incomplete penetrance of phenotype in the pdKO females can be attributed to a lack of recombination of one *Yap1* allele. Both *Progesterone receptor* and *Yap1* are located on chromosome 9 in mice with *Pgr* being located at 8899834-8968612 bp on the sense strand and *Yap1* being located at 7932000-8004597 bp on the antisense strand (Mgi). Both *Pgr* and *Yap1* are located at 2.46cM and due to the basics of recombination efficiency, the likelihood of recombination for these two genes is nearly 0%. Unfortunately, this contributes to the only possible genotype being *Pgr Cre* on the sense strand and one deleted allele of *Yap1* on the antisense strand within any given *Pgr* expressing cell within the murine uterus. The proximity of these genes led to the partiality of the double knockout and the lack of complete phenotype. Despite the incomplete penetrance of phenotype, we observed significant differences in pdKO females compared to controls, which compromised fertility. It is likely that complete conditional uterine knockout of *Yap1* combined with *Wwtr1* would lead to a much more severe effect, including fully compromised decidualization and fertility due to a lack of maternal remodeling. Currently, we are working to generate cell type-specific knockouts to determine the endometrial compartmental contributions of total *Yap1* and *Wwtr1* knockout.

## Materials and Methods

### Animal Models

*Pgr^Cre/+^*(19) mice were crossed to *Yap^fl/fl^ Wwtr1^fl/fl^*(20, 21) to generate *Pgr^Cre/+^ Yap^fl/+^ Wwtr1^fl/fl^* partial double knockouts (pdKO). The *Pgr^Cre/+^*mice are a mixed background of 129Sv × C57BL/6, the *Yap^fl/fl^ Wwtr1^fl/fl^* mice are 129SvEv. Animals were housed and maintained in a designated animal care facility at Michigan State University on a 12-hour light/dark cycle with free access to food and water. All animal procedures were approved by the Institutional Animal Care and Use Committee of Michigan State University.

### Fertility Evaluation

Females were co-housed with proven fertile wild-type males for a 6-month breeding trial. Males were rotated in or out of cages if females did not produce a live born litter within one-month from time of set up. For timed mating experiments, proven fertile wild-type males were placed in female cages in the evening. Seminal plugs were checked each morning with day of plug designated at 0.5 days post coitus (dpc). Tail vein injection with Chicago blue dye served as a positive identifier for implantation sites at all time points. Following blue dye injection, female mice were sacrificed for collection at a variety of timepoints including 1.5, 3.5, 4.5, 5.5, 7.5, 9.5, and 12.5dpc. Body weight, uterine wet weight, ovarian wet weight, implantation site number, and gross morphology were catalogued. Uterine, oviductal, and ovarian tissues were divided and flash frozen or stored in RNAlater for downstream RNA and protein analyses or fixed in 4% paraformaldehyde for histological analysis. A subset of tissues were fixed in 10% DMSO in methanol for tissue clearing and advanced light sheet microscopy.

### Artificial Decidualization

Sexually mature (8 weeks or older) female mice were ovariectomized followed by two weeks of rest. Animals were treated with three daily subcutaneous injections of 100ng 17β-estradiol followed by two days of rest then three daily injections of 1mg progesterone plus 6.7ng 17β-estradiol. Six hours following injection on the third day, an intraluminal scratch was performed surgically on the anti-mesometrial side of the endometrium of one uterine horn utilizing a blunted 25G needle. The unscratched uterine horn served as an unstimulated hormonal control. Decidualization reaction was maintained with daily subcutaneous injection of 1mg progesterone plus 6.7ng 17β-estradiol for 5 total days followed by euthanasia (n=5/genotype). Uterine tissues were collected as described above. Decidual reaction was measured by uterine wet weight ratio of stimulated/unstimulated horn and molecular markers *Wnt4* and *Bmp2* by qPCR of stimulated uterine horn compared to unstimulated horn from the same animal.

### Oviductal and Uterine Flushes

Sexually mature female mice (8 weeks or older) were mated to proven fertile wild-type males. Oviductal flushes were performed with phosphate buffered saline (PBS) by inserting a 30G needle into the infundibulum of both oviducts at 1.5dpc (n=6 per genotype). Flushed products were collected, counted, categorized, and imaged. Uterine and oviductal flushes were performed at 3.5dpc with uterine flushes performed by inserting a 30G needle into the uterotubal junction and flushing toward the cervix utilizing PBS. Flushed products were similarly collected, counted, categorized, and imaged.

### Immunohistochemistry

Tissues were fixed in 4% paraformaldehyde, dehydrated in ethanol and xylene, and embedded in paraffin. Sections (6 μm) were deparaffinized and rehydrated in a graded alcohol series followed by antigen retrieval (Vector Laboratories, Burlingame, CA) and hydrogen peroxide treatment. Next, sections were blocked and incubated with antibodies against YAP, WWTR1, ER-alpha, PGR, Ki67, SUSD2 or ASMA overnight at 4°C (see Supplementary Table 1 for IHC antibody information). On the following day, sections were incubated with biotinylated secondary antibodies followed by incubation with horseradish peroxidase conjugated streptavidin. Immunoreactivity was detected using the DAB substrate kit (Vector Laboratories) and visualized as brown staining by light microscopy. Incubation with secondary antibody only served as a negative control. Alternatively, after dehydration, slides were stained with Masson’s Trichrome followed by rehydration in a graded ethanol series then cover slipped and visualized by light microscopy. ImageJ image analysis software (NIH, v2.14.0), was utilized to determine a digital HSCORE for staining intensity of luminal epithelium, glandular epithelium, and stromal compartments of each uterine section or for granulosa cells in ovarian cross-sections.

### RNA isolation and real-time quantitative PCR

Total RNA was isolated from frozen mouse tissue using TRIzol reagent (Invitrogen, Waltham, MA). About 1μg of RNA was reverse transcribed to cDNA using a High-Capacity cDNA Reverse Transcription kit according to the manufacturer’s instructions (Applied Biosystems, Foster City, CA). Quantitative real-time PCR (qPCR) was performed with SYBR Green PCR Master Mix (Applied Biosystems) using the BioRad CFX Opus 384 qPCR system utilizing primers targeting genes of interest (Supplementary Table 2). Expression was normalized to the average of *36b4* and *18s* per sample and fold change determined compared to time matched floxed controls (*Pgr^+/+^ Yap^fl/fl^ Wwtr1^fl/fl^*).

### RNA-sequencing and data analysis

Total RNA was isolated from flash frozen mouse uteri utilizing TRIzol reagent as described above. Following isolation, RNA was treated with a RNA Clean and Concentrator kit (Zymo) followed by DNAse treatment (TURBO DNA-free kit, Invitrogen) and stored in nuclease-free water at -80°C. Concentration was determined utilizing a Qubit RNA BR Assay Kit (Invitrogen). Samples were sent for RNA integrity analysis then subsequently sent to a sequencing facility for library preparation and sequencing (Michigan State University Genomics Core). Libraries were prepped with Illumina Stranded mRNA Library Prep kit (Illumina) and sequenced (paired end 150bp) on a NovaSeq 6000 Instrument (Illumina) to an average depth of 40 million read pairs per sample. Reads were quality trimmed, adapters were removed using TrimGalore (version 0.6.10), and quality-trimmed reads were assessed with MultiQC (version 1.7). Trimmed reads were mapped to Mus Musculus GRCm39.110 and gene counts quantified using STAR (version 2.6.0c). Model-based differential expression analysis was performed using edgeR-robust method (version 3.42.4) (22) in R. Genes with low counts per million (CPM) were removed using the filterByExpr function from edgeR. Multidimensional scaling plots, generated with the plotMDS function of edgeR, were used to verify group separation prior to statistical analysis. DEG were identified as FDR P value less than 0.05. Visualization of DEG was performed utilizing EnhancedVolcano (version 1.18.0) to generate a volcano plot. Counts per million of selected DEG were plotted using the box plot function of ggplot2 (version 3.4.4). Gene set enrichment and visualization was completed using the ShinyGO online tool (version 0.80) (23).

### Whole-mount immunofluorescence, 3D uterine imaging, and analysis

Uteri were dissected from 3.5 (n=2) and 5.5dpc (n=8) pdKO and control females. Whole mount immunofluorescent staining was performed as previously described(24). Briefly, following animal euthanasia, uterine samples were subsequently fixed in DMSO:methanol (1:4) and stored at -20°C. Then, samples were rehydrated in 1:1, Methanol:PBST (PBS + 1% Triton X-100) for 15 minutes then washed for 15 minutes in 100% PBST. Tissues were then incubated in a blocking solution of PBS, 1% Triton X-100, and 2% powdered milk for 2 hours at room temperature. Samples were stained with primary antibodies for rat anti-CDH1 (M108, Takara Biosciences), rabbit anti-cytokeratin 8 (MA5-14476, Invitrogen) and rabbit anti-FOXA2 (Abcam, ab108422) diluted at 1:500 in blocking solution for 7 nights at 4°C. Uterine samples were then washed in PBST for 15 minutes twice and 45 minutes four times then incubated with fluorescently conjugated Alexa Fluor 555 Donkey anti-Rabbit IgG secondary antibody (A31572, Invitrogen), and 647 Goat anti-Rat secondary antibody (A21247, Invitrogen) and Hoechst (B2261, Sigma Aldrich) at 1:500 for 2 nights at 4°C. Samples were then washed in PBST for 15 minutes twice and 45 minutes four times, dehydrated in methanol then incubated overnight in 3% H2O2 diluted in methanol. Tissues were then washed in 100% methanol for 15 minutes twice then for 60 minutes and cleared overnight using BABB (benzyl alcohol:benzyl benzoate, 1:2). Stained tissue samples were imaged with a Leica TCS SP8 X Confocal Laser Scanning Microscope System (Leica Microsystems) with a white-light laser, using a 10x air objective. For each uterine horn, z-stacks were generated with a 7.0µm increment, and image analysis was carried out using Imaris v9.2.1 (Bitplane, Zurich, Switzerland). Briefly, confocal LIF files were imported into the Surpass mode of Imaris, and Surface module 3D renderings were used to create structures for the oviductal–uterine junctions, embryos, implantation chambers, and horns as described previously(25). We used the Contour modules for Hoechst and FOXA2 fluorescent signal (embryo), and CDH1 signal (oviduct and implantation chamber). Embryo 3D volume was assessed using the surface statistics function, while embryo implantation chamber length was calculated using the measurement points module of Imaris software.

### Statistical analysis

Data were tested for normality then log transformed in the case of non-normal data and analyzed by unpaired t-tests, one-way ANOVA, Fisher’s exact test, or mixed effects analysis as indicated. Data that were not normal were analyzed utilizing Kruskal-Wallis nonparametric tests. Statistical analyses were performed utilizing GraphPad Prism 10 (GraphPad Software) and values were considered significant if *p*<0.05.

### Data Availability

Raw FASTQ files were deposited in the NCBI Gene Expression Omnibus as GSE267798.

## LIST OF FIGURES AND LEGENDS

**Supplementary Figure 1. Analysis of ovaries from pdKO mice A.** mRNA analysis of *Yap1, Wwtr1, Ccn1,* and *Ccn2* of whole ovaries collected at 3.5dpc. **B.** DNA expression of floxed allelles in pdKO and control ovaries at 3.5dpc. **C.** Immunohistochemical analysis of YAP1 and WWTR1 in ovaries at 3.5dpc and semiquantitative HSCORE.

**Supplementary Figure 2. pdKO histoarchitechture is comparable to controls in primigravid and multigravida females. A.** Representative micrographs of trichrome staining of ovaries, oviducts, and uteri of virgin females at proestrus (8 weeks of age). **B.** Representative trichrome micrographs of ovaries, oviducts, and uteri at 3.5dpc in primigravid females (8-12 weeks of age). **C.** Representative micrographs of trichrome staining of ovaries (left panel), oviducts (center panel), and uteri (right panel) from multigravida breeding trial females (8 months of age).

**Supplementary Figure 3. pdKOs exhibit increased inter-litter timing and smaller litter sizes. A.** Expanded dataset showing decreased pups per litter in pdKOs compared to controls. **B.** pdKOs exhibit increased inter-litter timing. **C.** pdKOs exhibit a proportional body weight increase to controls over an 8-week period.

**Supplementary Figure 4. 3D imaging reveals normal oviductal fold patterning in pdKOs. A.** Representative micrographs of fluorescent whole mount imaging of oviducts of floxed controls (top panel) and pdKOs (bottom panel) at 3.5dpc in primigravid females. Images are whole oviduct (left), longitudinal folds (left middle), transverse folds (right middle), and continuous longitudinal folds (right) (white=Hoechst, blue=E-Cadherin, n=4). **B.** Representative micrographs of fluorescent imaging of embryos (marked with adjacent asterisk) within uteri at 3.5dpc (white=Hoechst, E-cadherin=blue). **C.** Embryos counted in uterine horns at 3.5dpc with whole mount fluorescent uterine imaging (n=2 females/genotype).

**Supplementary Figure 5. Estrogen receptor signaling is not aberrant in pdKOs. A.** *Esr1* and target genes, *Muc1, C3, Muc4* mRNA expression in 3.5dpc uteri. **B.** Immunohistochemical analysis of Estrogen receptor expression and semiquantitative HSCORE of positive DAB stain in 3.5dpc uteri.

**Supplementary Table 1. Antibodies used for western blot and immunohistochemistry analyses.**

**Supplementary Table 2. Primer sequences.**

**Supplementary Table 3. Breeding trial data.**

**Supplementary Table 4. Gene Ontology Enrichment for Biological Process for upregulated DEG.**

**Supplementary Table 5. Gene Ontology Enrichment for Biological Process for downregulated DEG.**

**Supplementary Table 6. Gene Ontology Enrichment for Cellular Component for upregulated DEG.**

**Supplementary Table 7. Gene Ontology Enrichment for Cellular Component for downregulated DEG.**

## Acknowledgements

The authors thank Dr. John Lydon (Baylor College of Medicine) and Dr. Francesco DeMayo (NIEHS) for the *Pgr Cre* mouse model, and Dr. Amy Ralston (MSU) for the *Yap* and *Wwtr1* floxed mouse models. The authors thank and acknowledge that this work was made possible by the excellent technical assistance of Samantha Hrbek, Erin Vegter, and the MSU Vivarium Staff. Research reported in this manuscript is supported by the Eunice Kennedy Shriver National Institute of Child Health & Human Development of the National Institutes of Health under award numbers R01HD099090 to A.T.F., and F31HD107923 to G.E.M. Training for G.E.M. was supported in part by the Eunice Kennedy Shriver National Institute of Child Health & Human Development of the National Institutes of Health under Award Number T32HD087166. Partial support for this work was also provided by MSU AgBio Research, and Michigan State University. This content is solely the responsibility of the authors and does not necessarily represent the official views of the National institutes of Health.

## Contributions

GEM, NJ, and ATF conceptualized the study. GEM designed experiments. GEM, NM, EV, and ILPD executed experiments. GEM, NM, and RA performed analyses. GWB, BG, RA, and ATF contributed to intellectual oversight, analytical guidance, and training of GEM. GEM wrote the manuscript. ATF contributed funding for this study. All authors reviewed and accepted the final version of the manuscript.

