## Supplementary Figures for "Yes Associated Transcriptional Regulator 1 (YAP1) and WW Domain Containing Transcription Regulator (WWTR1) are required for murine pregnancy initiation"

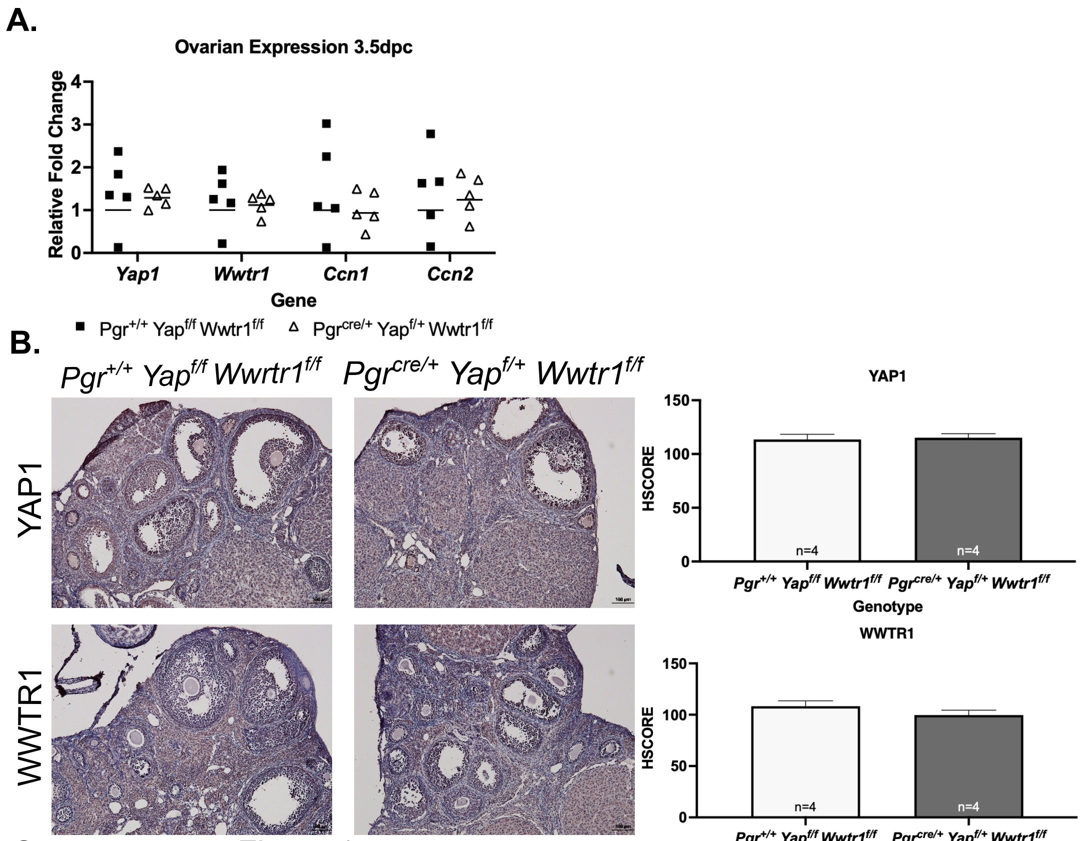


Supplementary Figure 1.

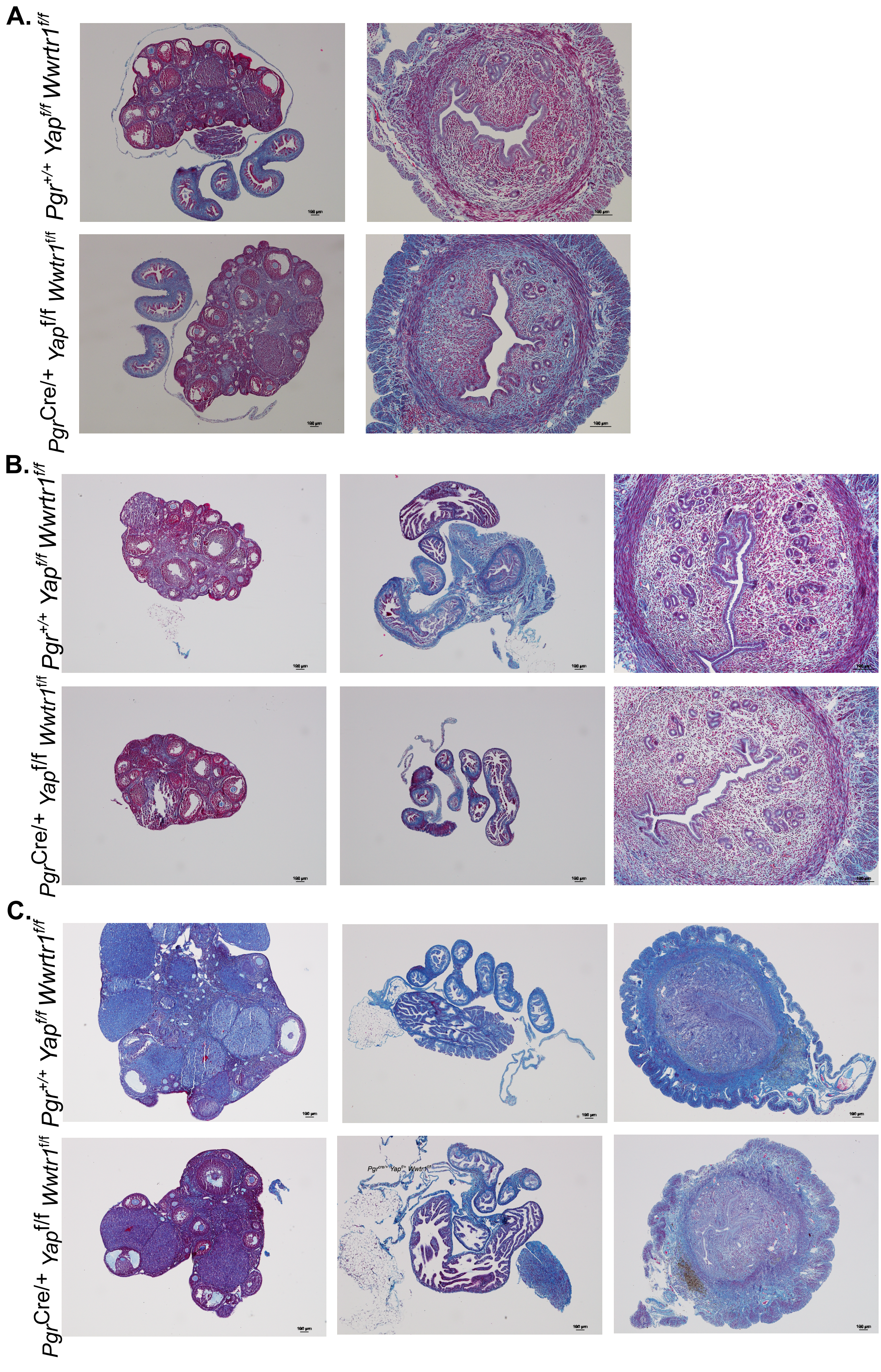


Supplementary Figure 2.

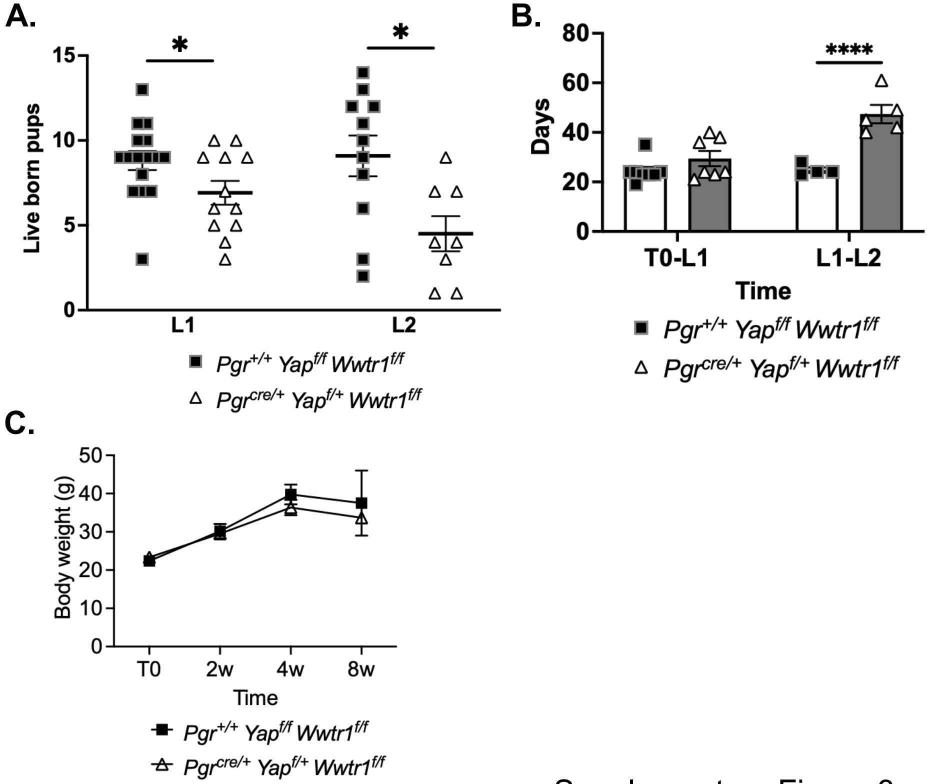


Supplementary Figure 3.

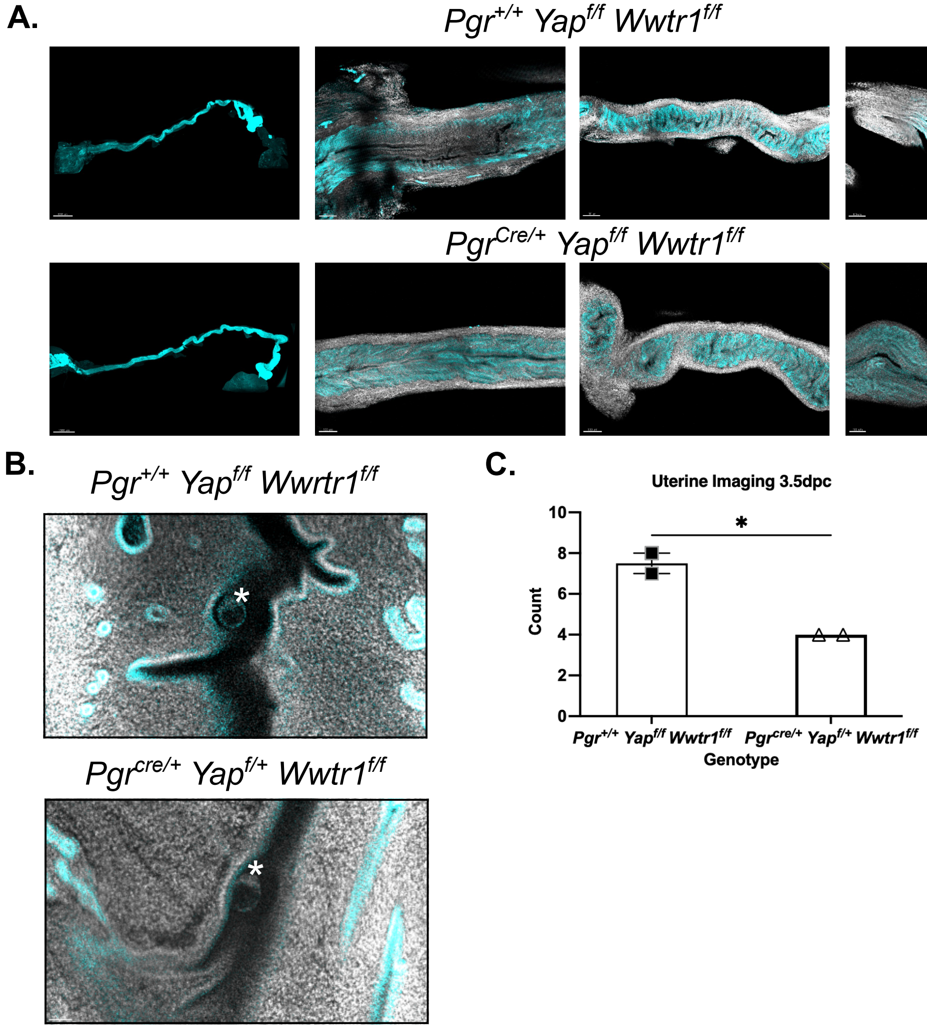


Supplementary Figure 4.

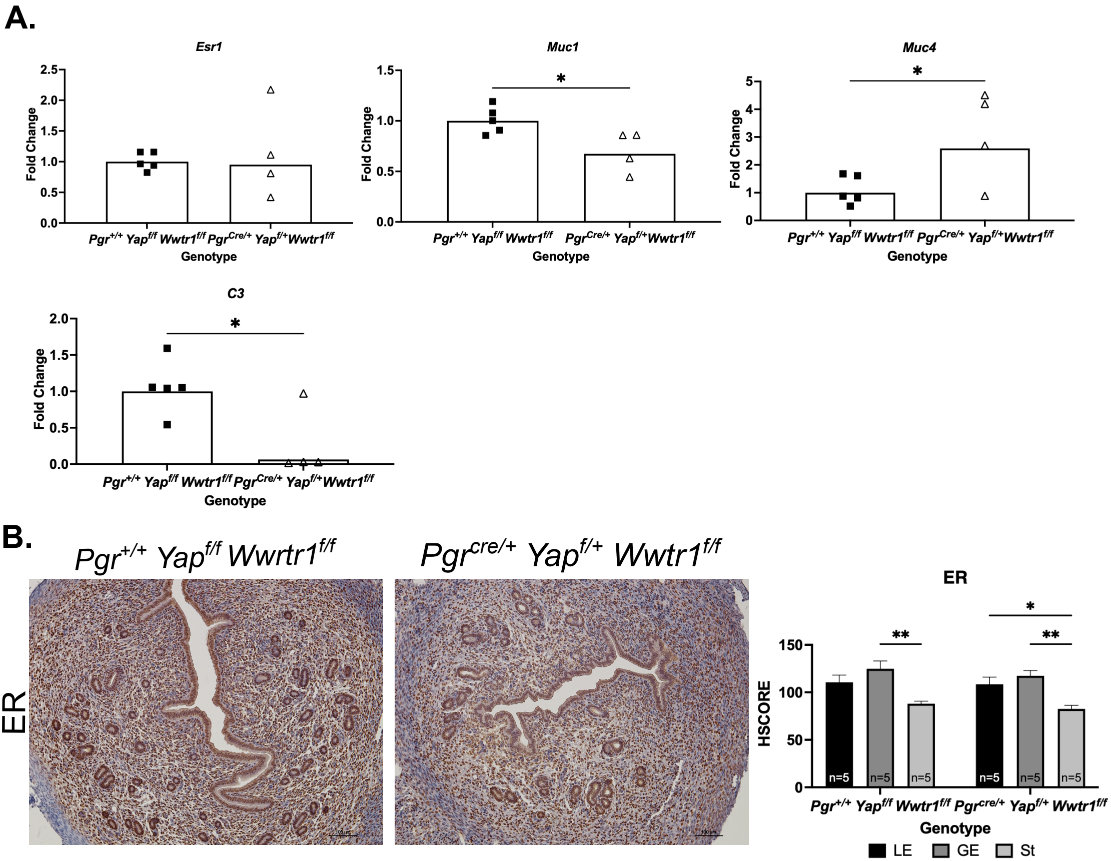


Supplementary Figure 5.
